# Nuclear exosome targeting complexes modulate cohesin binding and enhancer-promoter interactions in 3D

**DOI:** 10.1101/2025.08.25.671287

**Authors:** Charbel Akkawi, Alexandre Heurteau, Xavier Contreras, Marion Helsmoortel, Stéphane Schaak, Julie Bossuyt, Olivier Fosseprez, David Dépierre, Olivier Cuvier, Rosemary Kiernan

## Abstract

Three-dimensional long-range contacts between enhancers and promoters are thought to be largely determined by loop extrusion driven by the cohesin complex and insulator factors. However, recent evidence also suggests a role for noncoding RNAs (ncRNAs), such as enhancer-associated RNAs (eRNAs) and promoter upstream transcripts (PROMPTs), in shaping enhancer-promoter connectivity. While nuclear RNA exosome, together with targeting complexes, PAXT and NEXT, control the decay of ncRNAs, it has not yet been determined whether these complexes regulate 3D contacts. Chromatin recruitment maps of ZCCHC8 (NEXT), ZFC3H1 (PAXT) and MTR4 helicase revealed that these factors that associate with sites of enhancer-promoter interactions. Depletion of NEXT, PAXT or MTR4 induced the accumulation of ncRNAs, notably enhancer-associated RNAs (eRNAs) and promoter upstream transcripts (PROMPTs). Strikingly, this further increased cohesin levels at sites accumulating ncRNAs. Chromatin conformation capture analysis revealed that MTR4 modulates the 3D long-range contacts between enhancers with their distant TSS targets. Upon loss of MTR4, contacts at anchor points increase while intraloop contacts decrease, suggesting that MTR4 facilitates loop extrusion. These data highlight a key interplay between cohesin-mediated enhancer-promoter interactions and the regulation of ncRNAs by nuclear RNA exosome that is consistent with a role for RNA in genome folding.

## Introduction

The human genome needs to be highly compacted in order to occupy a nucleus that measures approximately 10 μm in diameter. Compaction must occur in a manner that permits coordinated gene expression. To achieve this, the genome is organized at multiple scales. Chromosomes are sub-divided into distinct active and inactive (A/B, respectively) compartments, and more locally folded into topologically associating domains (TADS) that correspond to regions favoring high frequencies of contacts between distant sites, such as enhancers and their target promoters, that lie within the same TAD. The establishment of TADs and loop structures is thought to be determined by the coordinated action of CCCTC-binding factor (CTCF) and the cohesin complex, a ring-shaped multimeric structure. TADs are thought to be formed via ‘loop extrusion’, a mechanism in which the cohesin ring entraps chromatin and extrudes loops, until blocked at TAD borders by CTCF or other insulating factors ^1–8^.

While the organization of the genome in 3D has been implicated in the regulation of gene expression, it is less clear how gene expression affects genome folding. Recent evidence suggests that RNAPII and its RNA product may indeed influence 3D genome interactions^9^. Several subunits of the cohesin complex, as well as CTCF, bind RNA in addition to DNA. Deletion of the RNA-binding domain of CTCF abolished about 50% of chromatin loops in mESCs^10,11^. Notably, TAD boundaries often express ncRNAs, which stabilize cohesin-CTCF interactions or facilitate CTCF recruitment. RNA-guided recruitment of CTCF has been shown to increase the efficiency of target search by several fold^12^. RNAPII was recently shown to be required for certain enhancer-promoter contacts and to antagonize loop extrusion^13,14^. Moreover, the recent development of 3D interaction maps of RNA has revealed interactions between enhancer-associated RNAs (eRNAs) and promoter proximal transcripts (PROMPTs) that parallel the functional connectivity of enhancers and promoters^15^. Mechanisms that control the abundance of such RNAs is likely to affect RNA-RNA spatial interactions and, consequently, genome looping and gene expression.

Many ncRNAs, including enhancer-associated RNAs (eRNAs) and promoter proximal transcripts (PROMPTs) are intrinsically unstable and are degraded by the nuclear RNA exosome, a multi-subunit barrel-shaped structure containing a helicase, MTR4 and 3’ to 5’ exoribonucleases, Dis3 and Exosc10^16^. RNA substrates must first be delivered to the nuclear exosome, which is carried out by two major targeting complexes: the Nuclear Exosome Targeting Complex (NEXT)^17^ and the polyA tail exosome targeting connection (PAXT)^18^. NEXT consists of a zinc-finger protein, ZCCHC8, together with RBM7 and the shared RNA helicase, MTR4. NEXT targets mostly short RNAs that are not polyadenylated. The second targeting complex, PAXT, is comprised of a zinc-finger protein, ZFC3H1, together with RBM26/RBM27, ZC3H3 and PABPN1, and targets polyadenylated RNAs^18,19^. We recently identified an additional PAXT subunit, polyA polymerase gamma (PAPγ), further strengthening the link between PAXT and polyadenylated RNA substrates^20^. RNAs targeted by PAXT and/or NEXT include not only aberrant transcripts, but many RNAs that are associated with gene regulatory regions, such as enhancer RNAs (eRNAs) and promoter-proximal transcripts (PROMPTs), also known as upstream antisense transcripts (uaRNAs)^19^. While nuclear exosome and its targeting complexes have been well characterized, whether the degradation of certain targets, such as eRNAs but also PROMPTs and ncRNAs, might affect genome organization and gene expression is not clear.

We recently described the genome-wide mapping of PAXT subunits, ZFC3H1, RBM26 and PAPγ^20^. Here, we have identified the genome-wide localization of ZCCHC8 subunit of NEXT, as well as the shared helicase, MTR4. Chromatin immunoprecipitation and high throughput sequencing (ChIP-seq) mapping revealed that NEXT subunit, ZCCHC8 and PAXT subunit, ZFC3H1, were detected primarily at transcription start sites (TSSs) but also at active enhancers. Consistent with their localization, loss of these factors was associated with stabilization of PROMPTs and eRNAs, as expected. Loss of MTR4 also strongly stabilized ncRNAs including eRNAs and PROMPTs, as described previously^17–19^. However, ChIP-seq mapping revealed that MTR4 was present at active enhancers, but was notably absent at TSSs, suggesting that its interaction with TSSs may be transient. By integrating ChIP-seq, RNA-seq and chromatin conformation capture (Hi-C) data, our data show that MTR4 contacts TSSs via 3D interactions between enhancers and promoters. Hi-C and ChIP-seq data furthermore showed that MTR4 modulates cohesin-dependent 3D contacts. Loss of MTR4 increases enhancer-promoter contacts at the anchor site but antagonizes loop extrusion. These data suggest that nuclear RNA surveillance factors contribute to chromatin loop formation and help shape genome architecture.

## Results

### PAXT, NEXT and MTR4 are recruited to specific sites on chromatin

We have previously shown that PAXT subunits, ZFC3H1, RBM27 and PAPγ, are associated with chromatin, frequently at TSSs that generate PROMPTs^20^. To gain further insight into nuclear exosome-dependent regulation of RNAs and ncRNAs, which may involve its recruitment to sites of RNA synthesis, we mapped the genomic localization of NEXT and the shared helicase, MTR4, by chromatin immunoprecipitation combined with high throughput sequencing (ChIP-seq). While MTR4, ZFC3H1 and ZCCHC8 recruitment sites frequently co-localized, they also showed some notable differences (Fig. 1a and Supplementary Fig. 1). Only a partial overlap between ZFC3H1 and ZCCHC8 recruitment sites was expected, which is consistent with their function in targeting specific RNAs^17–19^ (Fig 1b-d, Supplementary Fig. 1). MTR4 recruitment sites overlapped with 1251 and 627 recruitment sites of ZCCHC8 and ZFC3H1, respectively (Fig. 1b; Fisher’s exact test: p-values < 1e-126and <1e-235, respectively), consistent with the functions of PAXT or NEXT with MTR4 in targeting specific RNAs^17–19^. Surprisingly, however, the majority of MTR4 sites did not overlap with those of either ZFC3H1 or ZCCHC8 even though MTR4 is a subunit of both PAXT and NEXT macromolecular complexes. Further inspection using averaged profile analysis of ChIP-seq peaks showed that while recruitment of ZCCHC8, ZFC3H1 and MTR4 partially overlapped, ZFC3H1 and MTR4, in particular, showed some specificity in recruitment site preference (Fig. 1c-e, top panels). Ranking of ChIP-seq signal of ZCCHC8 or ZFC3H1 showed relatively moderate levels of MTR4 (Fig. 1c-d, bottom panels), though significantly above random signal as shown on average profile analysis (Fig. 1c-d, top panels). Similarly, averaged profile analysis and ranking of MTR4 ChIP-seq reads on heat maps further revealed only modest signal of either ZFC3H1 or ZCCHC8 (Fig. 1e, Supplementary Fig. S1C). These results suggest that while binding sites of ZFC3H1, ZCCHC8 and MTR4 overlap to some extent, many sites appeared to be independent.

**Fig. 1.**
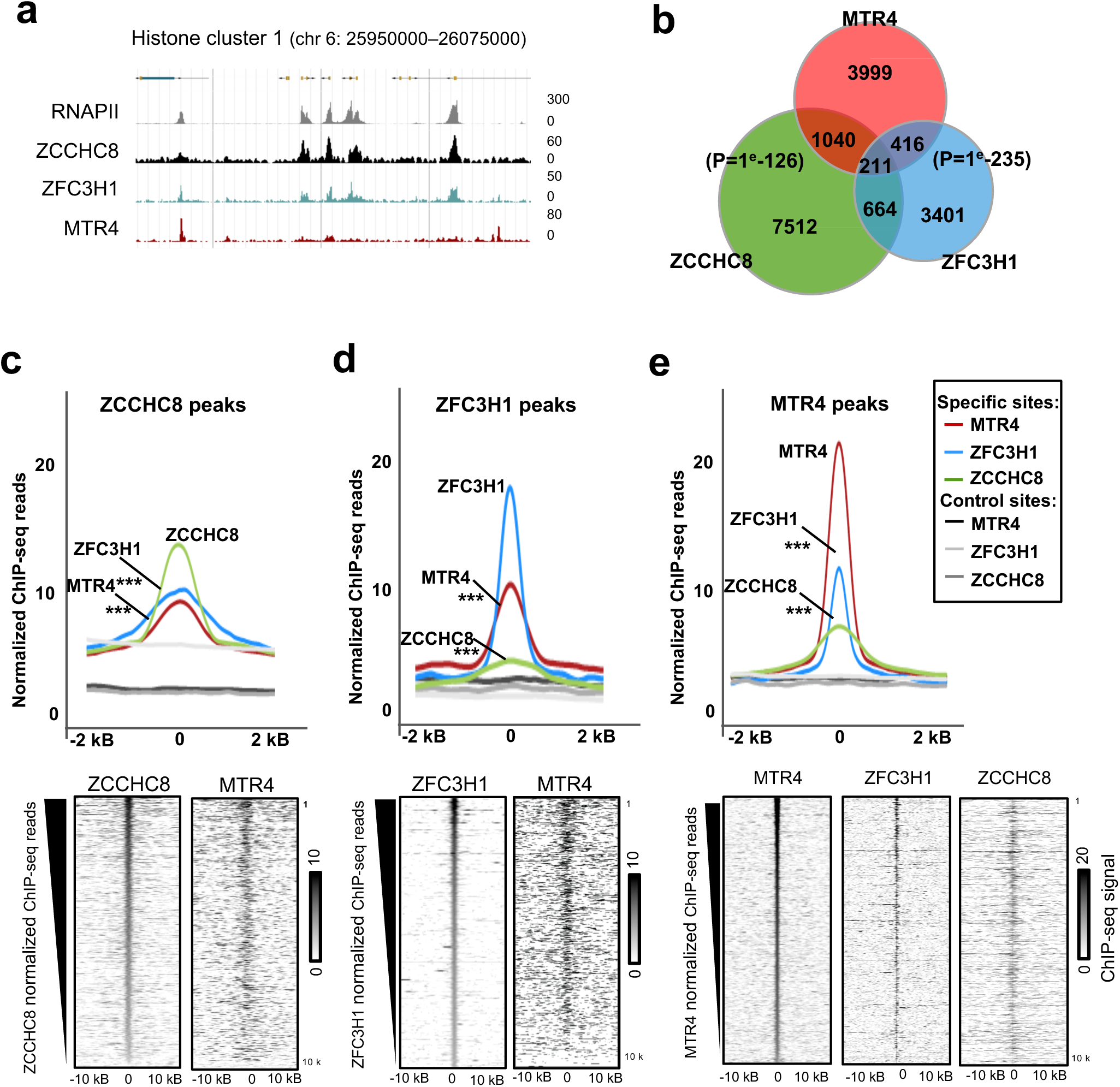
Genome-wide identification of sites bound by MTR4, ZFC3H1 or ZCCHC8. **a** Browsershot showing ChIP-seq reads following precipitation with anti-RNAPII, anti-ZCCHC8, anti-ZFC3H1 or anti-MTR4 specific antibodies, over the indicated region of the human genome. **b** Venn diagram showing the intersection between sites bound by MTR4, ZFC3H1 or ZCCHC8, as indicated. P-values were calculated by Fisher’s exact test. **c-e Top** Average plots of ChIP-seq reads of ZCCHC8, ZFC3H1 and MTR4, as indicated, at ZFC3H1, ZCCHC8 or MTR4 recruitment sites compared to control sites (see Methods). P-values were calculated using a Wilcoxon test for specific sites compared to control (random) sites. **bottom** Heatmaps centered and rank-ordered on normalized ChIP-seq reads of ZCCHC8, ZFC3H1 and MTR4, as indicated. ChIP-seq reads of ZCCHC8, ZFC3H1 or MTR4 were plotted respecting the same ranking, as indicated.

### Recruitment of PAXT, NEXT and MTR4 at ncRNA target sites

To further investigate binding profiles of NEXT, PAXT and MTR4, we compared sites of recruitment to those of synthesis of their ncRNA targets. To do so, we generated atlases for coding and non-coding RNA targets, using strand-specific RNA sequencing data obtained from cells depleted of MTR4 (Supplementary Fig. 2A), ZFC3H1 subunit of PAXT and ZCCHC8 subunit of NEXT^18,21,22^. As expected, depletion of MTR4, ZCCHC8 or ZFC3H1 led to the accumulation of thousands of ncRNAs (Supplementary Fig. 2B), using a high confidence threshold for detection (FDR p-value of 1e-6). The ncRNAs detected in each depletion condition showed significant overlap, with more than 500 ncRNAs detected upon depletion of either ZFC3H1 or ZCCHC8 that were also detected upon MTR4 depletion (Supplementary Fig. 2B; p-value of 1e-102 and 1e-149, respectively) and levels of >200 ncRNAs were increased upon depletion of ZFC3H1, ZCCH8 or MTR4 (Supplementary Fig. 2B; p-value of 1e-121).

The ncRNAs detected were classified into categories based on their localization with respect to genes (Supplementary Fig. 2C). A significant fraction of ncRNAs detected occur immediately upstream (<2 kb) from transcription start sites (TSSs) of coding genes (Supplementary Fig. 2C-D), corresponding to PROMPTs, in agreement with previous reports^18,19,21^. Approximately 30% of the ncRNAs increased upon depletion ZFC3H1, MTR4 or ZCCHC8 were identified as PROMPTs (Supplementary Fig. 2C). Other categories of ncRNAs increased upon depletion of ZFC3H1, ZCCHC8 or MTR4 include eRNAs, antisense RNAs within gene bodies, and RNAs in intergenic regions (Supplementary Fig. 2C). Among PROMPTs, 62.5% of ZFC3H1-dependent PROMPTs and 51.6% of ZCCHC8-dependent PROMPTs were also increased following MTR4 depletion (Supplementary Fig. 2E). Similarly, 54.6% of ZFC3H1-dependent eRNAs and 42.5% of ZCCHC8-dependent eRNAs were also increased following MTR4 depletion (Supplementary Fig. 2F). Our data are thus similar to previous analyses showing the contribution of MTR4-containing complexes in regulating ncRNAs including eRNAs and PROMPTs.

We then analyzed the genomic distribution of MTR4, ZFC3H1 and ZCCHC8 ChIP-seq reads with respect to their ncRNA targets. Surprisingly, less than 2% of MTR4 sites localized near a TSS, in contrast to ZFC3H1 sites that were more frequently associated with a TSS, and to a lesser extent ZCCHC8 sites (Fig. 2a). Accordingly, MTR4 recruitment sites accounted for < 5 % of the PROMPTs detected upon the depletion of ZFC3H1, ZCCHC8, or MTR4 itself (Fig. 2b; p-values of 1). In contrast, ZFC3H1 was detected at 27% of the PROMPTs detected upon its depletion (Fig. 2b; bottom left Venn diagram, p-value < 1e-11). Similarly, 28% of ZCCHC8-dependent PROMPTS co-localized with a ZCCHC8 recruitment site (Fig. 2B; bottom right Venn diagram). Consistent with the poor overlap between MTR4 binding sites and PROMPTs, recruitment of MTR4 was essentially not detected at TSSs bound by either ZFC3H1 or ZCCHC8 (Fig. 2c). Accordingly, the histone mark H3K4me3 that is typical of active TSSs was enriched at ZFC3H1 or ZCCHC8 sites, but not at MTR4 sites (Supplementary Fig. 2G). In contrast, MTR4 recruitment sites were enriched in the histone H3K4me1 mark that is associated with enhancers (Supplementary Fig. 2G). In support of this finding, ZFC3H1- and ZCCHC8-bound enhancers were associated with MTR4 (Fig. 2d). In summary, MTR4 co-localizes with ZFC3H1 or ZCCHC8 primarily at sites that bear marks of active enhancers, in stark contrast with TSSs where MTR4 recruitment is poorly detected.

**Fig. 2.**
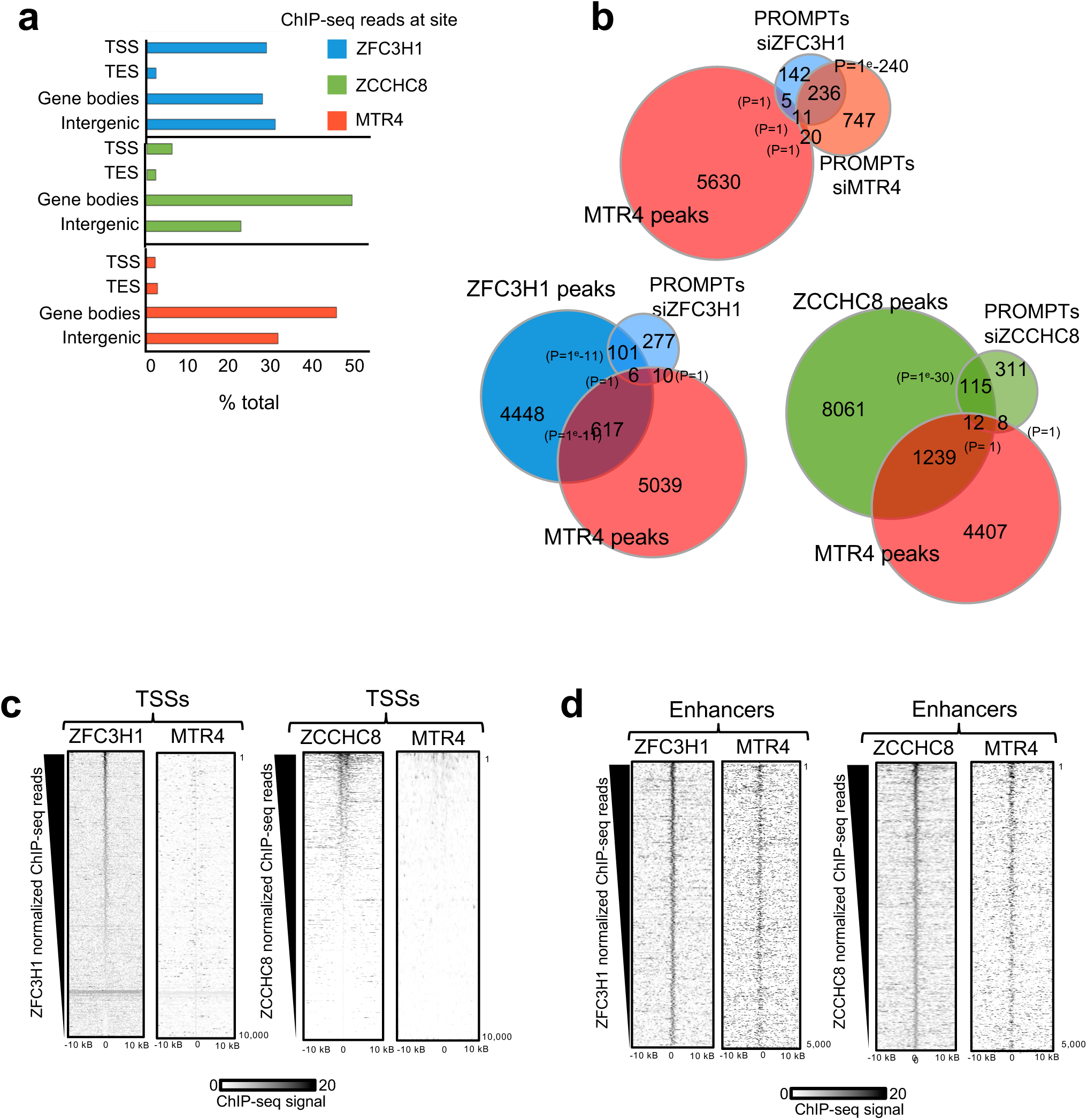
Recruitment sites of MTR4 overlap with those of ZFC3H1 and ZCCHC8 but not with PROMPTs. **a** Histograms showing the genomic distribution of ChIP-seq peaks of ZFC3H1, ZCCHC8 and MTR4, as indicated, overlapping with transcription start sites (TSSs), transcription end sites (TESs), gene bodies or intergenic (not surrounding gene units +/- 2kb) regions. **b** Venn diagrams showing the intersection between PROMPTs detected in cells depleted of MTR4, ZFC3H1 or ZCCHC8, as indicated, and binding sites of MTR4, ZFC3H1 or ZCCHC8, as indicated (P-values were obtained using Fisher’s exact test). **c** ChIP-seq heatmaps centered and rank-ordered by ZFC3H1 (left) or ZCCHC8 (right) signal at TSSs (+/- 10 kb). ChIP-seq reads of MTR4 were plotted respecting the same ranking, as indicated. **d** ChIP-seq heatmaps centered and rank-ordered by ZFC3H1 (left) or ZCCHC8 (right) signal at distant enhancer sites (+/- 10 kb). ChIP-seq reads of MTR4 were plotted respecting the same ranking, as indicated.

### Distal MTR4 sites establish long-range interactions with TSSs that generate PROMPTs

To understand how MTR4 may influence the accumulation of TSS-associated PROMPTs despite being poorly detectable at sites of PROMPT synthesis, we first determined the expression level of flanking genes in the presence and absence of MTR4, reasoning that increased expression of the flanking gene might explain the increase in PROMPTs. While the expression of genes flanking PROMPTs was sometimes deregulated in MTR4-depleted cells (126 genes) (Supplementary Fig. 3A), MTR4-dependent PROMPTs were associated with both up-regulated and down-regulated genes, similarly to ZFC3H1- or ZCCHC8-dependent PROMPTs (Supplementary Fig. 3B-E). This result argues against the possibility that the accumulation of MTR4-dependent PROMPTs was linked to increased expression of the flanking gene. Moreover, these results suggest that the differential expression of genes does not explain the effect of MTR4 on TSS-associated PROMPTs.

As MTR4 was recruited to distant sites marked by H3K4me1 (Supplementary Fig. 2G), our data raised an alternative possibility that MTR4, together with ZFC3H1 or ZCCHC8, may be involved in regulating PROMPTs depending on enhancer-promoter long-range interactions. Cohesin preferentially binds to H3K4me1-associated enhancers^23^ to mediate long-range interactions (LRIs) with distant TSSs that may be more or less transient^7,24^. Remarkably, more than half of all MTR4 binding sites co-localized with the Rad21 subunit of cohesin, and a significant proportion co-localized with both Rad21 and H3K4me1 (Fig. 3a and Supplementary Fig. 4A-B; Fisher exact test p-value of 1e-225 and 1e-152). Assessing long-range interactions in 3D by Chromatin Conformation Capture (Hi-C) using 2D interaction maps indicated that many recruitment sites of PAXT/NEXT/MTR4 established LRIs with distant TSSs that accumulate PROMPTs upon nuclear exosome depletion (Fig. 3b). To further test a possible role of LRIs in juxtaposing MTR4 to its TSS-associated target, we first performed a genome-wide ranking of all pairs of enhancers/TSSs from the highest to lowest interaction, based on normalized levels of Hi-C counts^25–27^ (Fig. 3c). TSSs were further scored for the presence or absence of PROMPTs. Of interest, PROMPTs were largely enriched at TSSs scoring high interaction levels with distal enhancers bound by MTR4 (Fig. 3c, bottom row of matrix). In contrast, TSSs having the least contact with distal MTR4 sites were significantly depleted of PROMPTs (Fig. 3c, bottom row). Moreover, TSSs with high levels of contacts with enhancers not bound by MTR4, were not significantly associated with PROMPTs (Fig. 3c, top row), indicating that stronger enhancer-promoter interactions alone do not explain the enrichment of PROMPTs. The differential levels of LRIs was validated by assessing the averaged physical contacts by aggregation of Hi-C data (aggregated plot analysis, APA), which readily estimates genome-wide LRIs^26,27^. APA showed that LRIs between enhancer/TSSs were clearly higher for the upper quartile that are enriched in MTR4-bound enhancers and PROMPTs compared to the lower quartile that are depleted of PROMPTs (Fig. 3d). These data suggest that PROMPTs are enriched at promoters engaged in high levels of LRIs with distal enhancer-bound MTR4. Moreover, these data fully agree with the formation of physical contacts between distant enhancers associated with nuclear exosome-associated sites and their target TSSs harboring PROMPTs (Fig. 3c, schematic, top panel). As such, these data raise the possibility that regulation of ncRNAs may involve the juxtaposition of enhancer-bound MTR4 complexes to distal TSSs, in 3D.

**Fig. 3.**
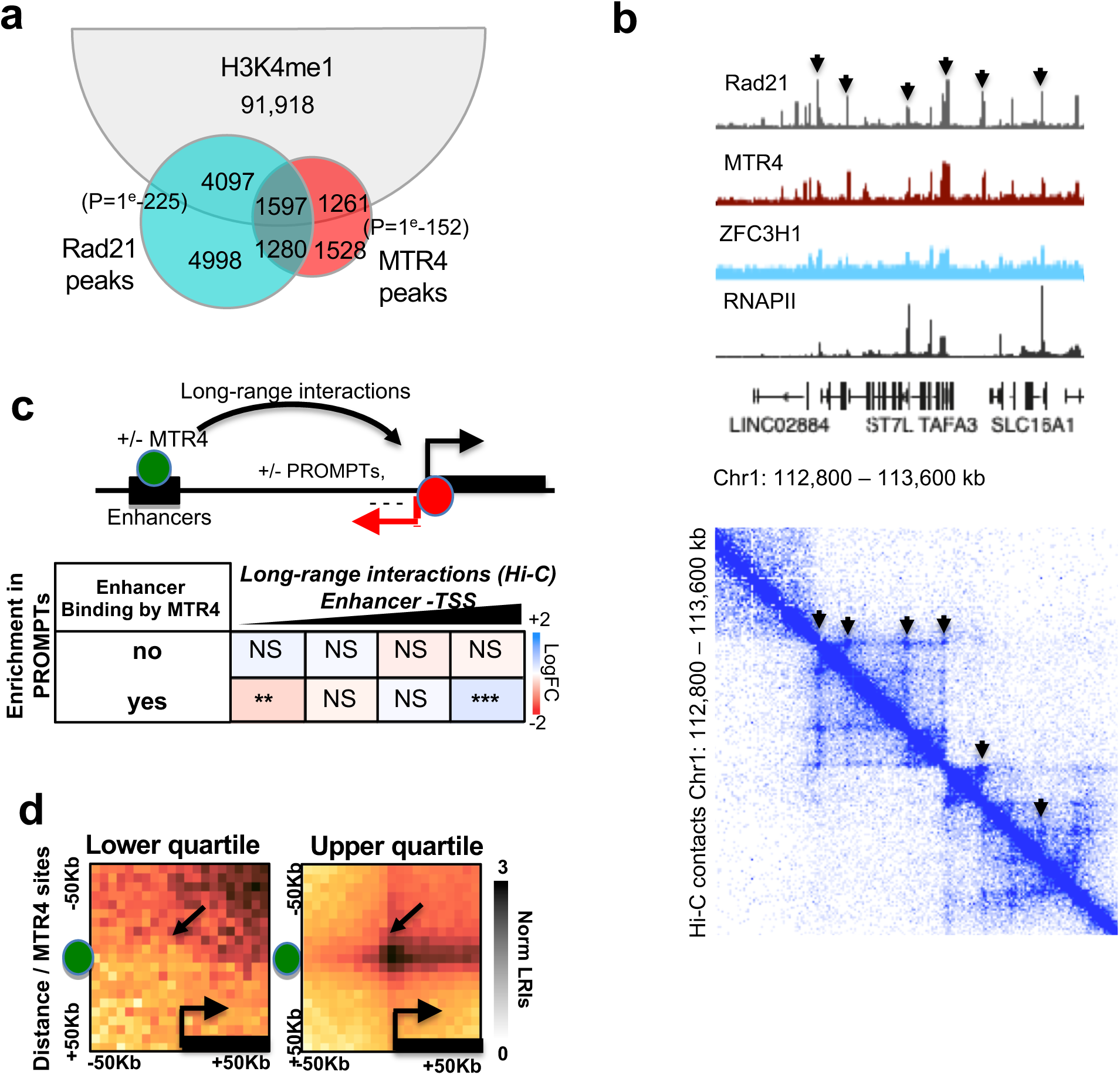
MTR4 influences PROMPTs through long-range interactions. **a** Venn diagram showing the intersection between binding sites of MTR4, Rad21 and the enhancer-associated H3K4me1 histone mark. P-values were obtained using Fisher’s exact test. **b** Top: Genomic view of the binding profiles of RNAPII, MTR4, ZFC3H1 and Rad21. Bottom: 2D matrix representing the levels of normalized long-range interactions within the region, determined from Hi-C data. **c** Top: Schematic representation of the enrichment test performed for enhancer-promoter LRIs, determined by systematic genome-wide interrogation of TSSs from Hi-C data, with the indicated features. Bottom: Matrices showing enrichment for the presence of PROMPTs, depending on levels of LRIs (ordered in quartiles) and co-localization of MTR4 with distant enhancers (**, p-value < 1e-5, ***, p-value < 1e-6, NS= not significant, Wilcoxon test). **d** 2D APA plots representing normalized Hi-C counts surrounding the MTR4 site (middle of the vertical y-axis) with sites surrounding TSSs (middle of the horizontal x-axis) from the lowest and highest quartiles of LRIs represented in c.

### MTR4 is associated with sites of cohesin-dependent long-range interactions and modulates cohesin binding

Since MTR4 co-localized significantly with the Rad21 subunit of cohesin and the H3K4Me1 mark associated with active enhancers (Fig. 3a and Supplementary Fig. 4A-B), we tested if MTR4 might be recruited to cohesin-associated LRIs genome-wide. Physical contacts were probed through aggregation of Hi-C data (APA). LRIs were further oriented in the direction of sense transcription, which allows an estimate of loop extrusion associated with the enhancer-promoter contacts. Hi-C data and MTR4 ChIP-seq data from HeLa cells were analyzed to determine the frequency of cohesin-associated LRIs and loop extrusion, associated with MTR4 co-localization or not (Fig. 4a). Firstly, the analysis revealed that MTR4 is present at distal enhancers that are engaged in 3D interactions with TSSs and that these LRIs are associated with loop extrusion (Fig. 4a, left panel). Although this phenomenon could be general, we noted that LRIs involving enhancers that were not associated with MTR4 (No MTR4) showed significantly less loop extrusion (Fig. 4a; right panel, Fig 4b). This is likely to be due at least in part to low Rad21 association at these sites, since MTR4 recruitment sites frequently co-localize with Rad21 (Fig. 3a and Supplementary Fig. 4A-B). Indeed, LRIs between Rad21-associated enhancers and TSSs showed strong loop extrusion while enhancers that were not associated with Rad21 showed significantly lower levels of loop extrusion, as expected (Supplementary Fig. 4C-D). Thus, MTR4 is associated with enhancers that engage in LRIs with TSSs and form extruded genome loops.

**Fig. 4.**
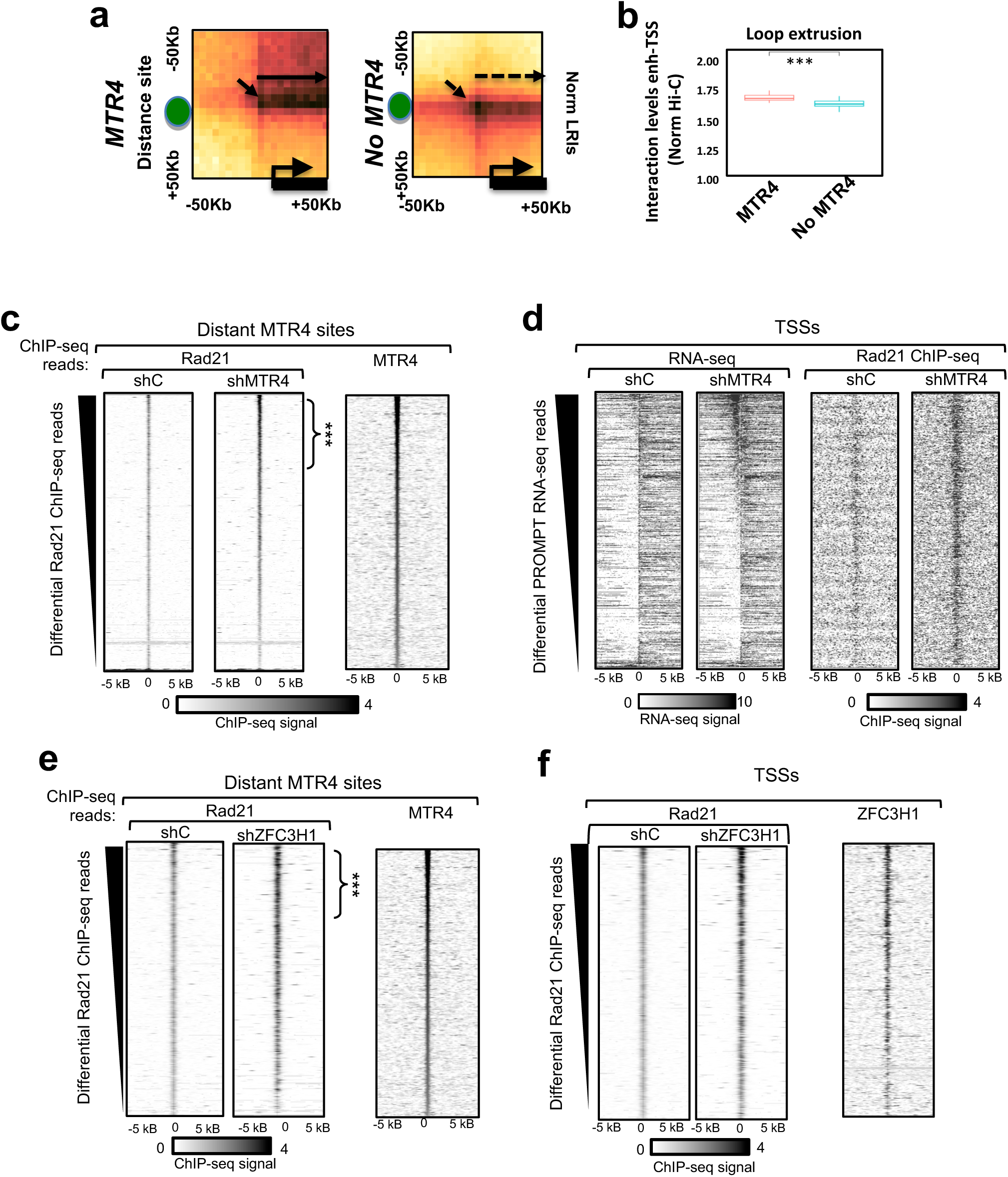
MTR4 is associated with sites of cohesin-dependent long-range interactions and modulates cohesin binding. **a** 2D APA plots representing normalized Hi-C counts at enhancers bound by MTR4 or not (left and right plot, respectively). Aggregation analysis was performed between enhancers and all active TSSs within 1 Mbp. **b** Box plot showing the distribution of long-range interactions along the bodies of genes contacted by distant enhancers depending on the presence or absence of MTR4. The statistical variations were tested by Wilcoxon test (*** p-value < 1e-6). **c** Heatmaps centred on MTR4 binding sites -/+ 5 kB, and rank ordered on differential Rad21 ChIP-seq reads in cells depleted of MTR4 (shMTR4) compared to control cells (shC) (left). MTR4 ChIP-seq reads were plotted respecting the same ranking (right) (*** p-value < 1e-6, Wilcoxon test). **d** Heatmaps rank ordered by increased PROMPTs detected by RNA-seq upon depletion of MTR4 compared to controls, and centred on TSSs -/+ 5 kB (left). Rad21 ChIP-seq reads were plotted respecting the same ranking. **e** Heatmaps centred on distant MTR4 binding sites -/+ 5 kB, and rank ordered on differential Rad21 ChIP-seq reads in cells depleted of ZFC3H1 (shZFC3H1) compared to control cells (shC) (left). MTR4 ChIP-seq reads were plotted respecting the same ranking (right) (*** p-value < 1e-6, Wilcoxon test). **f** Heatmaps of Rad21 upon depletion of ZFC3H1 compared to control, and centred on TSSs -/+ 5 kB (left). ZFC3H1 ChIP-seq reads were plotted respecting the same ranking (variation in Rad21 binding).

Since MTR4 frequently co-localizes with Rad21, we sought to test whether MTR4 might influence cohesin binding, as measured by ChIP-seq of cohesin in cells depleted of MTR4 compared to controls. To this end, we measured the association of cohesin with chromatin in cells depleted of MTR4 compared to controls by ChIP-seq of the Rad21 subunit. Strikingly, MTR4 depletion increased the accumulation of Rad21 that was bound at several hundred MTR4 sites (Fig. 5c and Supplementary Fig. 5A-B). Of note, Rad21 protein levels were unchanged in extracts of MTR4-depleted cells compared to controls (Supplementary Fig. 5C), indicating that the observed accumulation was not due to increased Rad21 expression. To determine whether such increases might be related to the abundance of ncRNAs, TSSs were then ranked by the accumulation of PROMPTs following MTR4 depletion (Fig. 4d, left heatmaps, highest to lowest PROMPT accumulation). Accumulation of PROMPTs at TSSs correlated strikingly with gain in Rad21 ChIP-seq signal in MTR4-depletion condition (Fig. 4d, right heatmaps). Furthermore, TSSs at which Rad21 most accumulated upon MTR4 depletion were also those at which PROMPTs increased upon depletion of ZFC3H1 or ZCCHC8, in addition to MTR4 (Supplementary Fig. 5D). We also observed that the most significant effect was associated with the differential expression of the flanking genes (Supplementary Fig. S5E), suggesting an association with gene expression. In addition, repeating the same analysis upon depletion of ZFC3H1 (Supplementary Fig. 5F) confirmed the impact on Rad21 signal (Fig. 4e-f), indicating that cohesin association is modulated by MTR4, likely in the context of PAXT. Accumulation of cohesin was similarly found over distant MTR4 sites and TSSs, depending on ZFC3H1 depletion. As observed for MTR4, depletion of ZFC3H1 did not alter the expression of Rad21 in cell extracts (Supplementary Fig. 5G). Therefore, our results show that nuclear RNA exosome, which regulates the abundance of non-coding RNAs, also modulates the accumulation of cohesin at the same sites.

**Fig. 5.**
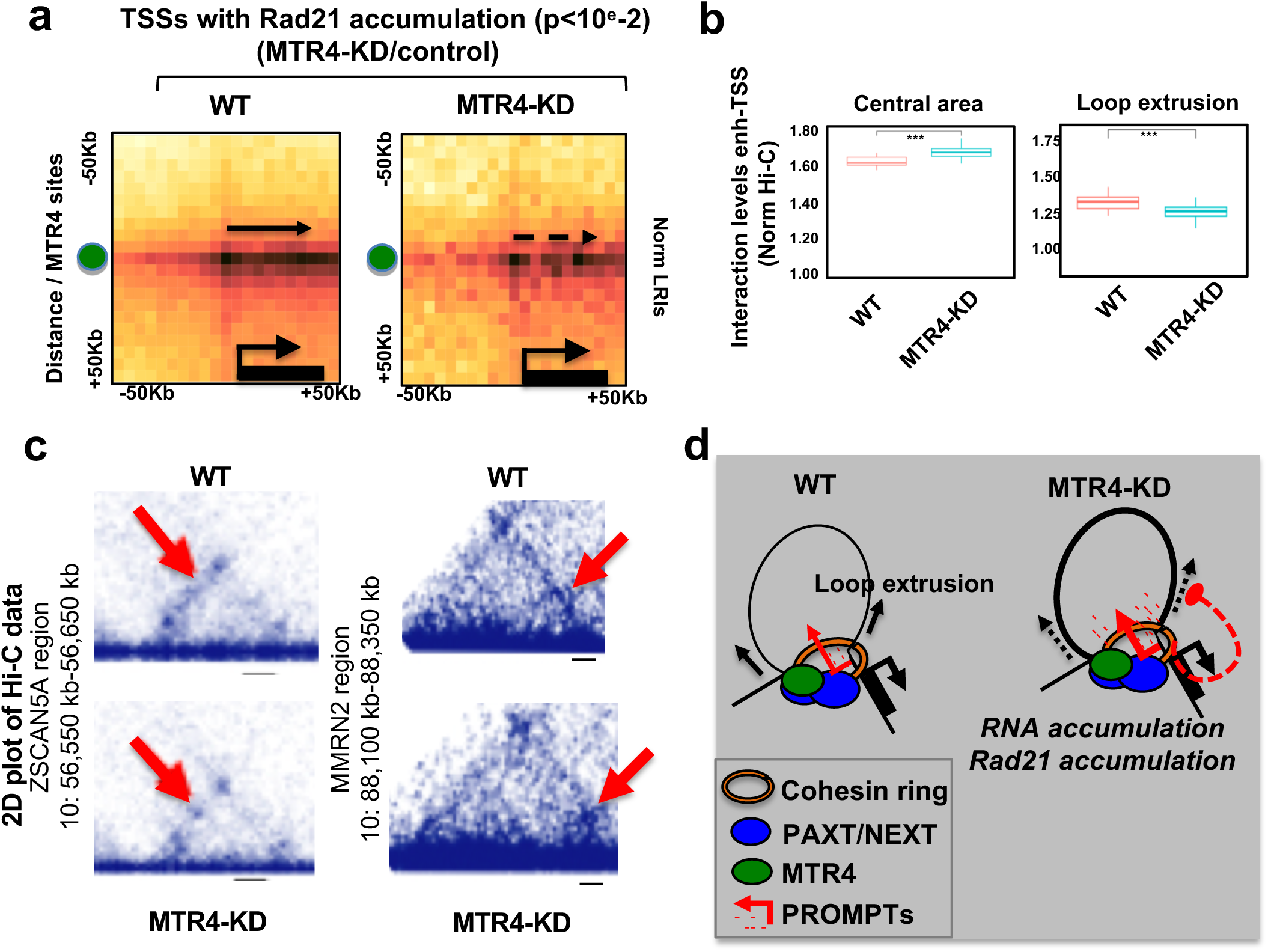
Nuclear exosome targeting complexes influence cohesin-mediated long-range interactions. **a** 2D APA plots representing normalized Hi-C counts at distant MTR4 sites in control or MTR4-depleted conditions (left and right plot, respectively). Aggregation analysis was performed between enhancers and all active TSSs within 1 Mbp. **b** Box plot showing the distribution of long-range interactions at interaction sites (central area, left box plot) or along the bodies of genes contacted by distant enhancers (loop extrusion, right box plot) in control or MTR4-depleted conditions, as indicated. The statistical variations were tested by Wilcoxon test (*** p-value < 1e-6). **c** Browser shots of Hi-C signal over representative regions in control and MTR4 knock-down cells, as indicated. Genomic coordinates in GRCh37 are shown on left. **d** Schematic representing a model where cohesin-mediated looping is influenced by PAXT, NEXT and MTR4. In wild-type cells (WT), ncRNAs including PROMPTs and eRNAs are degraded by nuclear exosome complexes found at loop contact sites. MTR4 depleted cells (MTR4-KD) show accumulation of eRNAs and PROMPTs and stabilization of cohesin-mediated loops (thick black line). Note that eRNAs may be still degraded at enhancers since MTR4 binds to enhancers with PAXT/NEXT, independently of cohesin-mediated looping, in contrast to MTR4-mediated degradation of PROMPTs at promoters that is associated with chromatin looping.

Since the accumulation of cohesin, that is involved in the formation of long-range contacts, was modulated by MTR4 at its recruitment sites, we wondered whether its depletion might also affect 3D genome interactions. Thus, aggregation of Hi-C data obtained from control cells and cells depleted of MTR4 was performed (Supplementary Fig. 6A). TSSs showing accumulation of Rad21 following loss of MTR4 (p <10^e^-2) were selected for analysis. Consistent with an increased association of Rad21, LRIs between TSSs and distant sites were significantly enhanced at the central region (Fig 5a and Fig. 5b, left panel). Of interest, contacts in the intra-loop region were significantly decreased following loss of MTR4 compared to the control condition (Fig. 5a and Fig. 5b, right panel). Interestingly, this pattern resembles that of LRI sites that are not associated with MTR4 in control conditions (Fig. 4a-b). Changes in LRIs were visible at specific MTR4-associated genomic regions (Fig. 5c). Taken together, these data show that LRIs are influenced by nuclear RNA exosome complexes containing MTR4.

## Discussion

Here, we identify an intricate regulation of 3D long-range interactions between enhancers and TSSs by nuclear RNA exosome complexes. We began by mapping the genomic localization of targeting complexes, PAXT and NEXT, via ZFC3H1 and ZCCHC8 subunits, respectively, and their shared RNA helicase subunit, MTR4. Comparison between the localizations of the complexes and ncRNA accumulated upon their depletions, revealed an interplay between the stability of ncRNAs, including PROMPTs and eRNAs, and 3D long-range interactions. Nuclear exosome subunits were found to be localized at sites that engage in long-range genomic interactions. Moreover, loss of nuclear exosome subunits, MTR4 or ZFC3H1, led to significant changes in the accumulation of cohesin, a key player in long-range interactions, at both TSSs and enhancers and modified the dynamics of chromatin looping (Fig. 5D).

Increasing evidence suggests that RNAPII and its RNA product may play a role in genome organization. Treatment with RNase abolished almost half of all genome contacts^10^. CTCF and cohesin subunits are RNA-binding proteins and mutation of the RNA binding region (RBR) of CTCF significantly disrupted CTCF-dependent insulation of chromatin domains and loop extrusion^10,11^. More recently, RNAPII has been shown to shape genome architecture by blocking loop extrusion while facilitating enhancer-promoter contacts^13,14^. Our data show that depletion of nuclear RNA exosome targeting complexes impacts 3D genome architecture and binding of cohesin. It seems unlikely that MTR4 or associated complexes, such as PAXT, are directly responsible for the effects observed on LRIs. Instead, it seems more probable that ncRNA targets of nuclear exosome, particularly eRNAs and PROMPTs, that were concomitantly stabilized at the enhancers and promoters in contact, may be responsible for the effect on LRIs and loop extrusion. Indeed, these data are consistent with recent studies showing that enhancer-promoter interactions are facilitated by RNAPII^13,14^ as well as pair-wise interactions between eRNAs and PROMPTs^15^. Thus, stabilization of ncRNAs, such as eRNAs and PROMPTs, following depletion of nuclear exosome targeting complexes would be expected to modify RNA-RNA spatial interactions and consequently, affect genome looping and gene expression.

Interestingly, our analysis revealed that although interactions between enhancers and TSSs and association with the Rad21 subunit of cohesin were increased, loop extrusion appeared to be less efficient. We speculate that insulator proteins, such as CTCF, Mediator^28^ or RNA-binding proteins, such as hnRNPK^15^, may be retained at the LRI sites, which may impede or contribute to a reduction in loop extrusion. The genome-wide mapping of nuclear exosome targeting complexes, described previously^20^ and in the present study reveal a strong association with TSS regions of actively transcribing genes and RNAPII. While still speculative, these findings raise the possibility that an important function of these complexes may be to fine-tune the abundance of TSS-associated and enhancer-associated RNAs and, consequently, modulate genome architecture.

An unexpected finding of genome-wide mapping was the absence of detectable MTR4 at TSSs, even though subunits of PAXT and NEXT are readily detectable at TSSs. MTR4 was more stably associated with enhancer regions and was present at sites engaged in long-range interactions with TSSs. This might suggest that cohesin-mediated LRIs may influence ncRNAs post-transcriptionally, depending on spatial targeting of the nuclear RNA exosome. Interestingly, Rahman et al. demonstrated that eRNA synthesis precedes looping and that their production is strongly reduced in the closed loop conformation^29^. Thus, chromatin loops might help degrade eRNAs when they are no longer required, by increasing nuclear exosome recruitment, or facilitating the activation of nuclear exosome locally, by juxtaposing the targeting complexes with MTR4 helicase whose binding is required to activate nuclear exosome. LRIs may thus define the context in which the nuclear exosome degrades ncRNAs, notably eRNAs and antisense PROMPTs. Improper accumulation of these ncRNAs may deregulate, either directly or indirectly, expression of the neighboring genes.

## Materials and Methods

### Cell culture and reagents

HeLa cells were grown in Dulbecco’s modified Eagle’s minimal essential medium (DMEM) (Sigma-Aldrich, D6429), supplemented with 10% fetal calf serum (FCS; Eurobio Scientific, CVFSVF00-01) and containing 1% penicillin-streptomycin (Sigma-Aldrich, P4333). HEK-293T were grown in Hepes-modified DMEM (Sigma-Aldrich, D6171), supplemented with 10% FCS (Eurobio Scientific, CVFSVF00-01) and containing 1% penicillin-streptomycin (Sigma-Aldrich, P4333). All cells were grown in a humidified incubator at 37°C with 5% CO2.

### Antibodies

Antibodies used in this study are shown in Table S1.

### RNAi

Production of short hairpin RNA (shRNA)-expressing lentiviral particles was performed as described previously (20) using plasmids expressing shRNAs targeting MTR4 (Sigma-Aldrich MISSION shRNA, TRCN0000296301), ZFC3H1 (Sigma-Aldrich MISSION shRNA, TRCN0000130498) or a non-targeting control (Addgene, plasmid 1864). For knockdown experiments, HeLa cells were either transduced with lentiviral particles and harvested 5 days later, or transfected with siRNAs (10 nM) shown in Table S2 using Interferin (PolyPlus) and harvested 72 h later, as described previously^30^. RNAi depletions were validated by immunoblot of total cell extracts, as described previously^30^.

### Sample preparation for RNA-seq, ChIP-seq and Hi-C

For RNA-seq, total RNA was extracted from HeLa cells using TRIzol (Thermo Fisher Scientific) according to the manufacturer’s instructions. RNA-seq (paired end, 125 bp) was carried out by BGI Genomic Services in triplicates.

Chromatin immunoprecipitation followed by high throughput sequencing (ChIP-seq) was performed from HeLa cells as described previously^31^ using the ChIP-IT High Sensitivity® kit from Active motif (ref #53040) according to the manufacturer’s instructions. Each ChIP used 30 μg of chromatin along with 4 μg of antibody detecting MTR4, ZFC3H1, ZCCHC8 or Rad21 (Supplementary Table S1). ChIP-seq libraries were constructed using the Next Gen DNA Library Kit (Active Motif 53216 and 53264). Library quality was assessed using Agilent 2100 Bioanalyzer and Agilent High Sensitivity DNA assay. High throughput sequencing (PE75) was performed by Sequencing-By-Synthesis technique using a NextSeq 500 (Illumina) at Genom’ic facility, Institut Cochin, Paris.

Hi-C was performed from HeLa cells using the Arima Hi-C ® kit from Arima Genomics (ref #A510008) according to the manufacturer’s instructions. Libraries were prepared using the KAPA® Hyper Prep kit (KK8500 and KK4824) and Illumina adapters (ref 20016329). High throughput sequencing (PE150) was performed by Novogene (NovaSeq) or BGI (DNBseq).

### RNA-seq alignment and ChIP-seq data integration

Raw RNA-seq data were aligned on hg19 reference genome using STAR aligner in paired-end and strand-aware mode. Resulting BAM files were used as such for differential analysis. For the detection of non-coding RNAs, all aligned BAM files were further split in two files corresponding to the strands of alignment.

After quality control using fastQC tool, ChIP-seq reads were aligned on hg19 reference genome using Burrows-Wheeler Aligner (BWA, http://bio-bwa.sourceforge.net/)^32^ and then normalized on the corresponding input as described previously ^33^. ChIP-Seq peak calling were performed in all conditions (depleted or not) with MACS2 with R 3.4.2 version. Gene TSSs associated with such peaks were identified by overlapping within the +/-500bp region around TSSs. Enhancers associated with peaks were identified by overlapping peaks summit with each enhancer range (+/-1000bp). Overlapping analyses were performed using ‘GenomicRanges’ package functions of R (https://bioconductor.org/packages/release/bioc/html/GenomicRanges.html). Venn Diagrams and statistical enrichment tests were performed using Fisher’s exact test or proportional tests. Boxplots showing quantification of ChIP-seq reads at the indicated genomic coordinates as performed using a Wilcoxon test comparing WT, MTR4-KD, ZFC3H1-KD or ZCCHC8-KD conditions, over the lists of gene loci. Boxplots were generated using ggplot2^34^ R package functions (https://cran.r-project.org/web/packages/ggplot2/index.html). Heatmaps were generated by aligning genes (oriented towards the right) on TSSs (position 0). Genes were ranked according to normalized ChIP-seq reads (TSS +/- 500 bp) of ZFC3H1 followed by visualization of the indicated ChIP-seq reads (of ZFC3H1, MTR4 or ZCCHC8), or of ncRNAs. Visualisation of heatmaps was performed with SeqPlot package^35^. H3K4me1, and Rad21 peaks on Hela cells were obtained from Encode Project (GEO accession GSM798322 and GSM935571 respectively).

### Detection of non-coding RNAs

Differential non-coding RNA levels were detected in MTR4 (this study), ZFC3H1 or ZCCHC8 depleted cells^18,21,22^. NcRNAs levels were compared between each condition and with control mock-depleted (‘WT control’) wild-type cells using enrichR function from NormR (binsize of 500 bp, FDR of 1e-5)^36^. Preliminary analyses were performed in comparison with analyses by mNET-seq and chromatin RNA-seq data in siLUC control and EXOSC3 siRNA conditions escribed previously^37,38^ .

### Heatmaps of RNA-seq and ChIP-seq data

Heatmaps were generated by aligning genes (oriented to the right) on TSSs (position 0) and by plotting the normalized bigwig signal, on reverse and forward strand for RNA-seq, simultaneously for each condition. Genes were ranked according to normalized ChIP-seq reads (TSS +/- 500 bp) of a given feature as indicated in the figure. Visualisation of heatmaps was performed with SeqPlot package^35^.

### Plots

Boxplot, Scatterplots, histograms and profiles were generated with ggpubr package based on ggplot2 package^34^. Venn diagrams were performed using Vennerable v1.0 package, developed by Jonathan Swinton.

### Computational integration of Hi-C data

Hi-C data at 5 Kb resolution were obtained from WT Hela cells or from cells depleted of MTR4. Hi-C data were validated by comparison with Hi-C in WT cells from Rao and colleagues (GSE63525)^25^ or Hi-C in wild-type and PDS5 depletion condition from J.M. Peter’s lab (GSE102884)^39^. Extraction of long-range contacts was performed using dump command from juicer tools in order to obtain normalized contacts (observed/expected and Knight-Ruiz for matrix balancing normalization) and filtered to fit inside Hela TADs. APA plots were generated as described ^24^ with heatmap function from R stats library after importing dumped matrices in 2D/3D, as previously described by ‘Aggregate Peak Analysis’ (APA)^27,40^, run on the normalized Hi-C matrices at a resolution of 10 kb as described^26,27,40^ with some adaptations. A minimal/maximal distance threshold of 200 kb to 1 Mbp was applied and each pair of sites was oriented to align the interactions between distant enhancers +/- bound by Cohesin/MTR4 sites and transcription start sites (TSSs) with their gene body towards the right of the APA. For statistical validation, a sampling of the interactions was performed to assess 50 independent times the interactions of 5,000 pairs in every condition. The variation was then tested using a Wilcoxon pairwise test for the central square values (3x3 central square (CS) of each of the 50 aggregated matrices) between the top and control group (e.g. top left corner of the APA matrix) applying a Wilcoxon test (stat_compare_mean from ggpubr, R package version 0.6.0). We estimated loop extrusion (all squares on the right/3’ that align with the central pixel) in parallel to the loop values (by taking the same number of pixels from the APA matrix (either 3x3 central square or 9 pixels on the right; each of the 50 aggregated matrices) and compared values using a Wilcoxon test (stat_compare_mean from ggpubr, R package version 0.6.0). Enrichment of a given site was performed by ranking all enhancer-TSSs interactions within a range of 200 kb to 1 Mbp according to their interactions strength (Normalized Hi-C counts) to define quartiles of enhancer-TSSs pairs from low-to-high (q1 to q4) interaction levels. Then, the proportion of a given feature (e.g. MTR4 sites) was scored for each quartile and enrichment was assessed using a Fisher’s exact test with respect to all combinations. Visualization of Hi-C signal was performed using Juicebox using default settings^41^.

### Statistical analyses of ChIP-Seq data

After quality control using fastQC tool, ChIP-seq reads of MTR4, ZFC3H1, ZCCHC8 and Rad21 and were aligned on hg19 reference genome using Burrows-Wheeler Aligner (BWA, http://bio-bwa.sourceforge.net/)^32^ and then normalized using the corresponding input. ChIP-Seq peaks were identified using MACS2 with normalization to the corresponding input sequenced in parallel generated in R. All subsequent analyses were done with R 3.4.2 version. Genes associated with peaks were identified by overlapping within the +/-500bp region around TSSs as defined by human TxdB database based on hg19 annotations (UCSC). Enhancers were identified by CAGE-seq (Encode). Overlapping analyses were performed using « GenomicRanges » R functions (https://bioconductor.org/packages/release/bioc/html/GenomicRanges.html), as described previously^33^. Enrichments in Venn diagrams and enrichment matrices were performed using Fisher’s exact test, p-values are either one-sided or two-sided to reflect enrichment and under-representation (i.e. positive Odds ratio, alternative = « greater » of R Fisher Test). Proportional Venn diagrams were plotted with « Vennerable » R package (https://github.com/js229/Vennerable). Strand-specific normalized read quantifications were performed by aligning stranded reads over oriented TSSs (+/- 5000bp). For analyses of enhancer-promoter associations, enhancers were those defined by Cap analysis of gene expression (CAGE-seq) by the Functional annotation of the mammalian genome (Fantom5) and Encode projects^42,43^.

### Statistical analyses of gene expression data

Gene expression analyses were first performed by polyA+ RNAseq analysis from HeLa cells depleted of MTR4 (MTR4-KD), ZFC3H1 (ZFC3H1-KD) as compared to siRNA treated control cells (‘WT’) in HeLa cells. Normalized strand-specific RNAseq reads were visualized using the IGV Gbrowser. Normalized strand-specific reads were further quantified in duplicates and directly plotted onto the heatmaps depending on gene ranking according to the indicated levels of ChIP-seq reads. After filtering low reads genes using R package HTSfilter, a differential analysis was performed with DESeq2 R package for comparing expression level of each gene in MTR4-KD or ZFC3H1-KD compared to siRNA treated WT conditions. Up-regulated and down-regulated genes subsets were defined according to a p-value threshold <0,05 and a LogFoldChange threshold > 1.5 or < 1.5 respectively. Enrichment tests between differentially expressed genes and other subsets of genes, such as those associated with peaks, were done using Fisher’s Exact test. Intersection matrices were performed using GRange with Fisher’s exact test.

## Data availability

RNA-seq, ChIP-seq and Hi-C data have been deposited at GEO (GSM4263550, GSE154100 and GSE283061). All data are available in the main text or the supplementary materials.

## Acknowledgements

The work carried out in this study was supported by ERC CoG ‘RNAmedTGS’, MSD Avenir ‘EpiMum3D’, Sidaction, LabUM EpiGenMed, to RK, MSD Avenir ‘Hit-Hidden-HBV’ to OC, and ANR ‘Helico’ and ANRS (#229035) to OC and RK. CA was supported by a fellowship from ANRS and DD by scholarships from La Ligue Nationale Contre le Cancer (LNCC) and ANRS.

We wish to thank members of the Kiernan and Cuvier labs as well as Matthias Merkenschlager for helpful discussions.

## Author contributions

XC and MH prepared ChIP-seq samples; CA performed Hi-C, CA and JB performed cell culture, RNAi and immunoblotting; AH, DD, SS, OF and OC performed data analysis; OC and RK conceived and supervised the project; OC and RK wrote the paper with input from all authors.

## Competing Interests

The authors declare no competing interests

## Additional Information

Correspondence and requests for materials should be addressed to Olivier Cuvier or Rosemary Kiernan.

## Notes

### Competing Interest Statement

The authors have declared no competing interest.

