## supplementary figs and tables for "Nuclear exosome targeting complexes modulate cohesin binding and enhancer-promoter interactions in 3D"

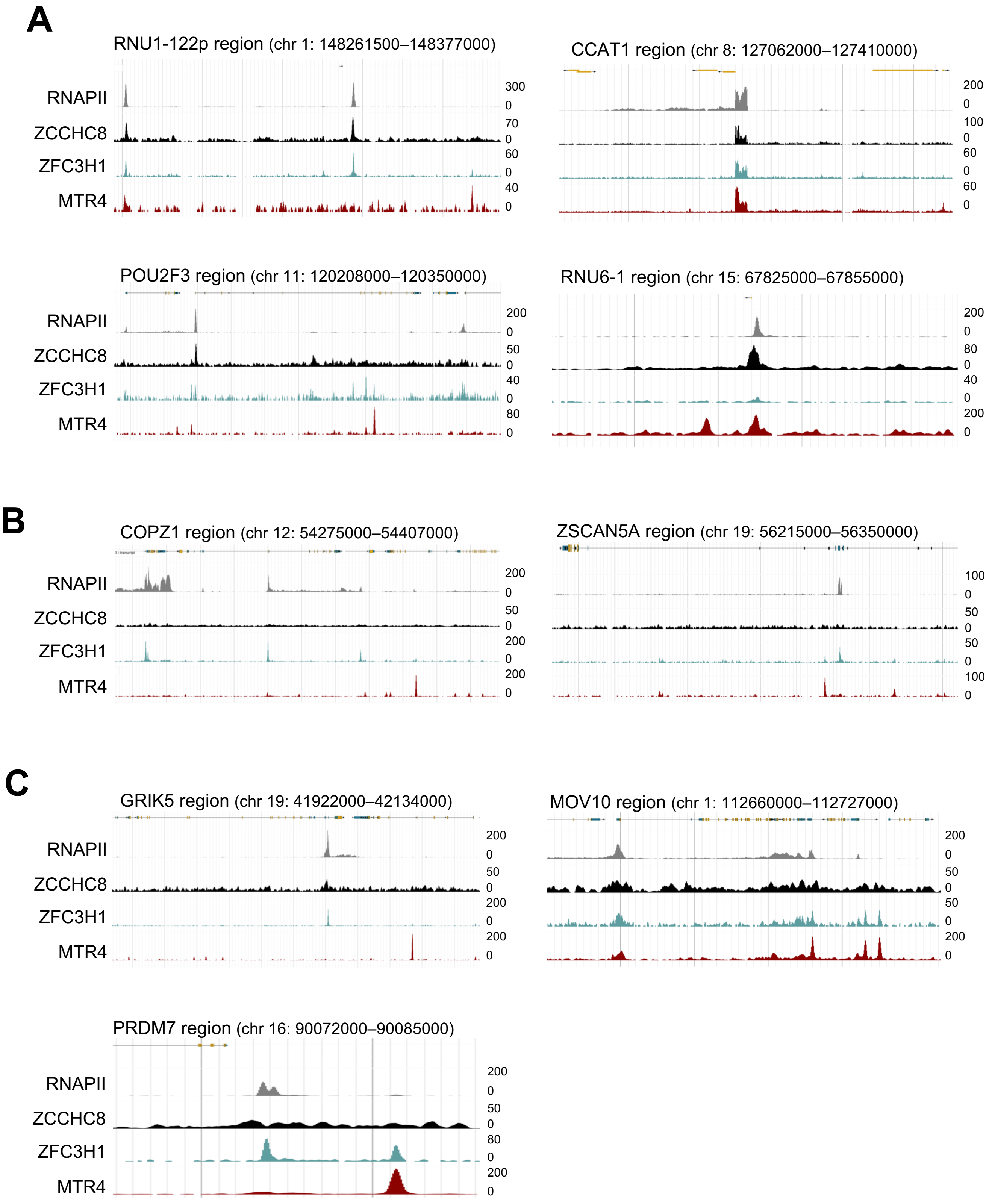


**Supplementary Fig. 1**

**Supplementary Fig. 1. Genome-wide localization of PAXT, NEXT and MTR4. A-C** Browser shots of RNAPII, ZCCHC8, ZFC3H1 and MTR4 ChIP-seq signal over representative genes in HeLa cells. A schematic representation of the gene is shown above. Shown are representative regions highly associated with ZCCHC8 (**A)**, ZFC3H1 (**B)** or MTR4 (**C)**.

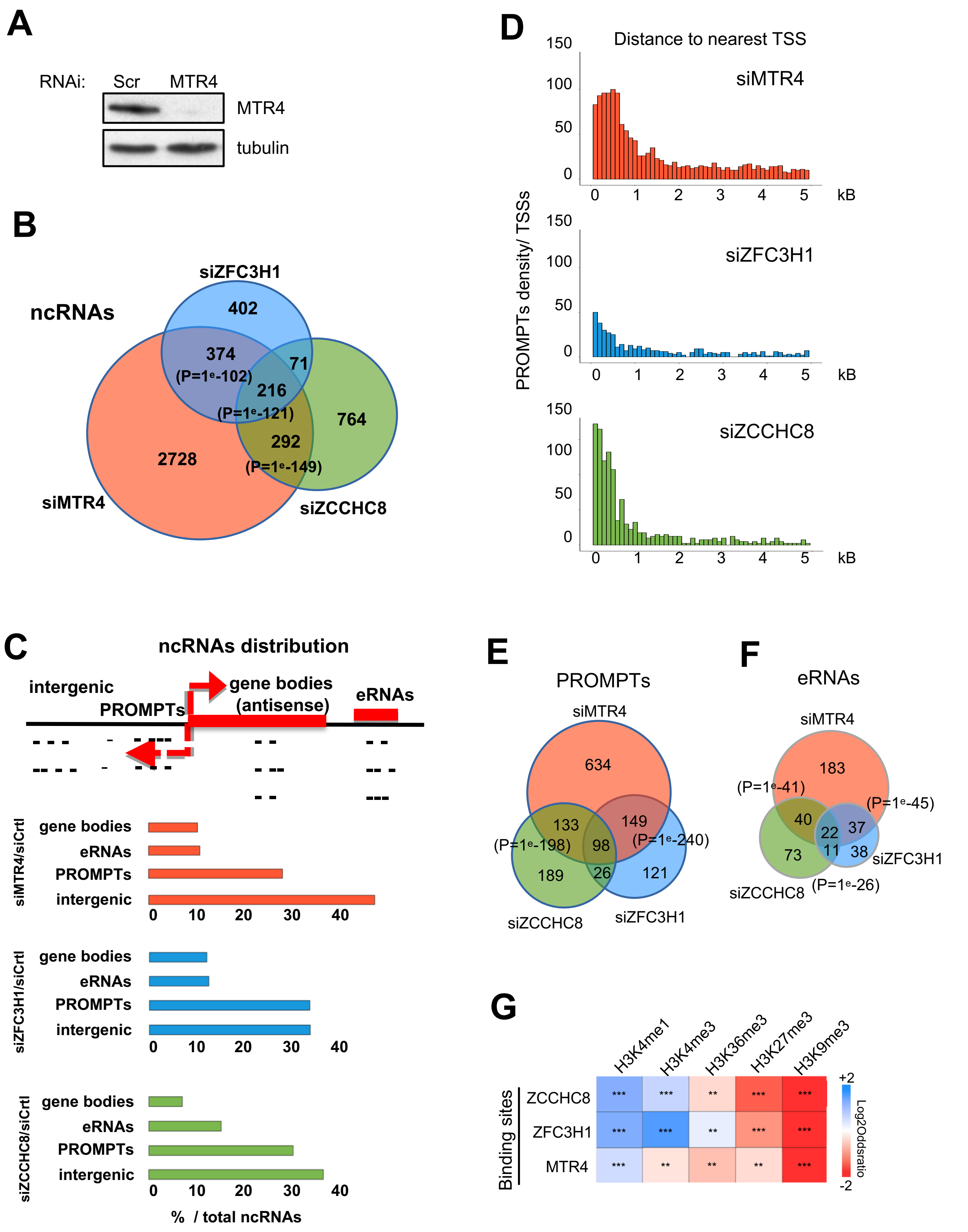


**Supplementary Fig. 2**

**Supplementary Fig. 2. Genome-wide detection of non-coding RNAs and Promoter upstream transcripts (PROMPTs) in cells depleted of MTR4, ZFC3H1 or ZCCHC8.**

**A** Whole cell extracts from HeLa cells treated with siRNAs targeting MTR4 or a non-targeting control, as indicated, were analysed by Western blot using the indicated antibodies. **B** Venn diagram showing the intersection between ncRNAs whose abundance increased in cells depleted of MTR4, ZFC3H1 or ZCCHC8 compared to control-depleted cells. P-values were calculated using Fisher's exact test. **C** (top) Schematic representation of categories of ncRNAs detected by differential expression of RNA-seq. (bottom) Histograms showing the distribution of ncRNAs detected upon depletion of MTR4, ZFC3H1 or ZCCHC8 as indicated, corresponding to gene bodies (antisense transcripts), enhancers (eRNAs), PROMPTs (antisense transcripts localized within 0 to 2 kb upstream of transcription start sites (TSSs)) and intergenic RNAs (ncRNAs present in regions away from -2kb to + 2kb from 5' and 3' end of genes, respectively). **D** Graphical representation of the distribution of the distance between ncRNA reads increased upon depletion of MTR4, ZFC3H1, or ZCCHC8 and the nearest TSS. **E** Venn diagram showing the intersection between PROMPTs whose abundance increased in cells depleted of MTR4, ZFC3H1 or ZCCHC8 compared to control cells. P-values were calculated using Fisher's exact test. **F** Venn diagram showing the intersection among eRNAs, detected in cells depleted of MTR4, ZFC3H1 or ZCCHC8 compared with control depleted cells. P-values were calculated using Fisher's exact test. **G** Intersection matrix showing the relative enrichments of the binding sites of MTR4, ZFC3H1 and ZCCHC8 with those of post-transcriptional modifications of histones associated with enhancers (H3K4me1), promoters/TSSs (H3K4me3), gene bodies (H3K36me3) or heterochromatin (H3K27me3 and H3K9me3). P-values were calculated by Fisher's exact test; log-ratios are indicated by color scale.

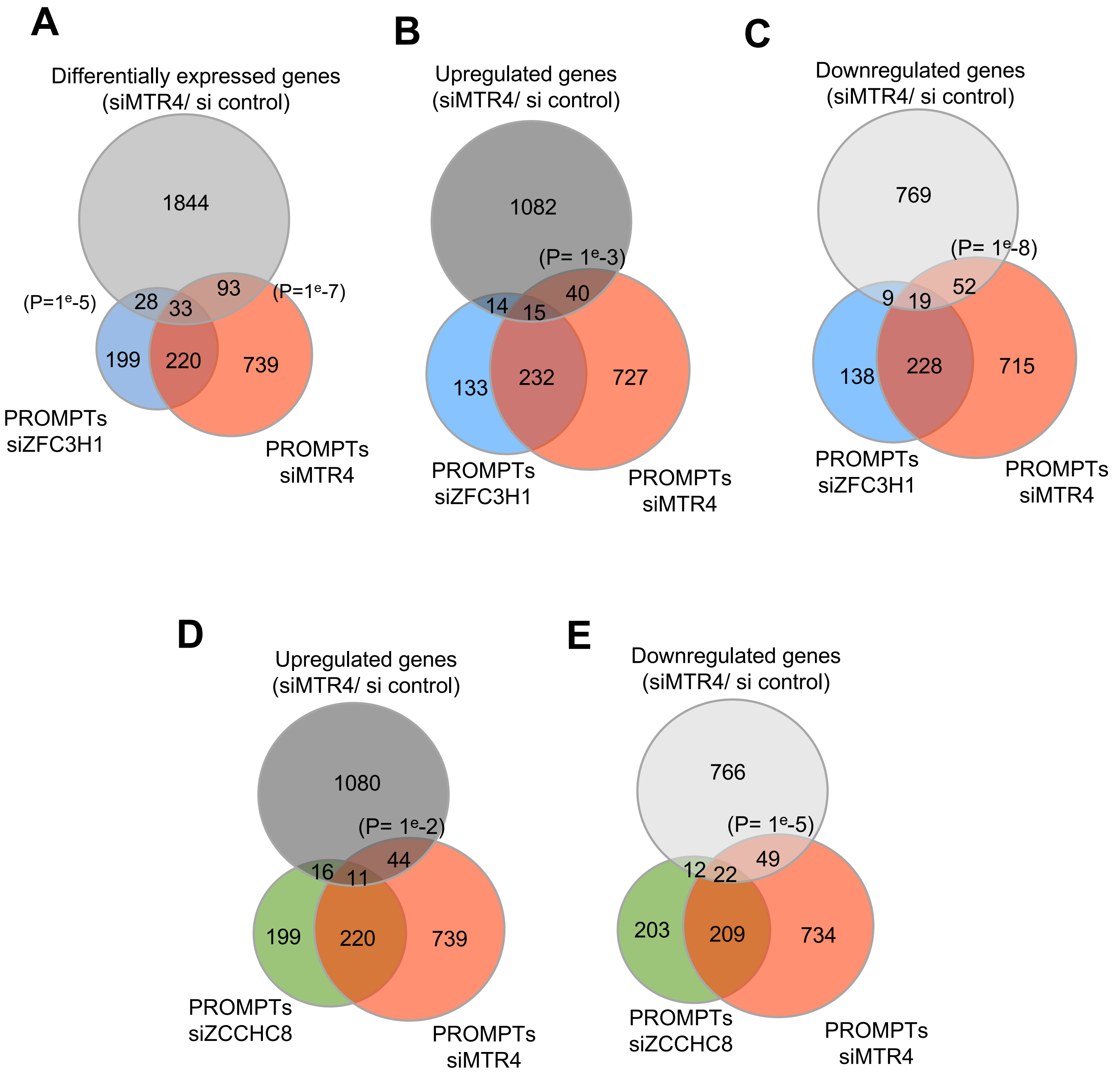


**Supplementary Fig. 3**

**Supplementary Fig. 3. MTR4 influences PROMPTs through long-range interactions.**

**A** Venn diagram showing the intersection between PROMPTs detected in cells depleted of MTR4 or ZFC3H1, as indicated, and the most differentially expressed genes upon depletion of MTR4. P-values were obtained using Fisher's exact test. **B** Venn diagram showing the intersection between PROMPTs detected in cells depleted of MTR4 or ZFC3H1, as indicated, and genes whose expression increased upon depletion of MTR4. P-values were obtained using Fisher's exact test. **C** Venn diagram showing the intersection between PROMPTs detected in cells depleted of MTR4 or ZFC3H1, as indicated, and genes whose expression decreased upon depletion of MTR4. P-values were obtained using Fisher's exact test. **D** Venn diagram showing the intersection between PROMPTs detected in cells depleted of MTR4 or ZCCHC8, as indicated, and genes whose expression increased upon depletion of MTR4. P-values were obtained using Fisher's exact test. **E** Venn diagram showing the intersection between PROMPTs detected in cells depleted of MTR4 or ZCCHC8, as indicated, and genes whose expression decreased upon depletion of MTR4. P-values were obtained using Fisher's exact test.

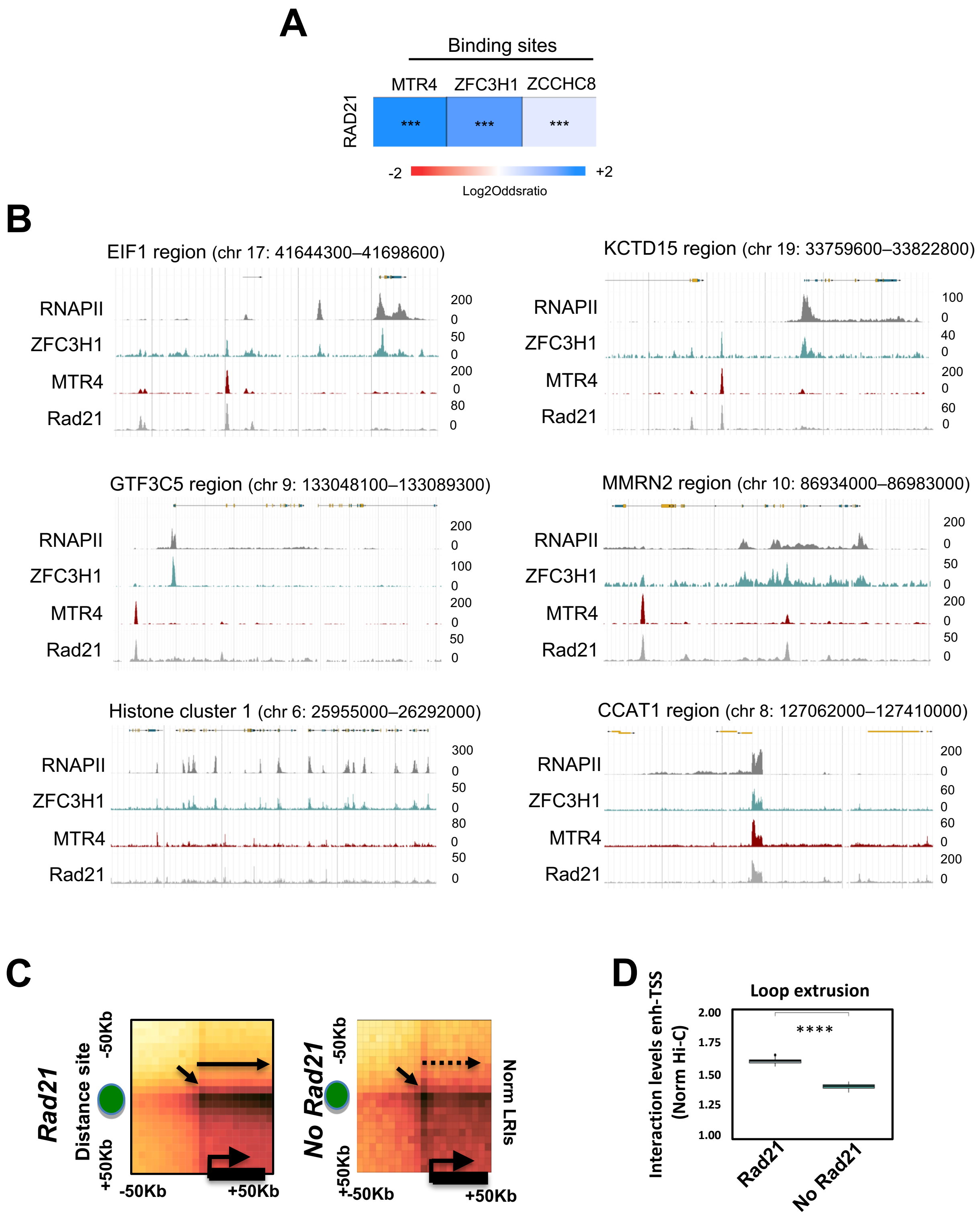


**Supplementary Fig. 4**

**Supplementary Fig. 4. MTR4 colocalizes with cohesin.**

**A** Intersection matrix showing the relative enrichments of the binding sites of MTR4, ZFC3H1 and ZCCHC8 with those of Rad21. P-values were calculated by Fisher's exact test; log-ratios are indicated by color scale. **B** Browser shots of RNAPII, ZFC3H1, MTR4 and Rad21 ChIP-seq signal over representative regions in HeLa cells. A schematic representation of the region is shown above. **C** 2D APA plots representing normalized Hi-C counts at enhancers bound by Rad 21 or not (left and right plot, respectively). Aggregation analysis was performed between enhancers and all active TSSs within 1 Mbp. **D** Box plot showing the distribution of long-range interactions along the bodies of genes contacted by distant enhancers depending on the presence or absence of Rad21. The statistical variations were tested by Wilcoxon test (**** p-value < 1e-6).

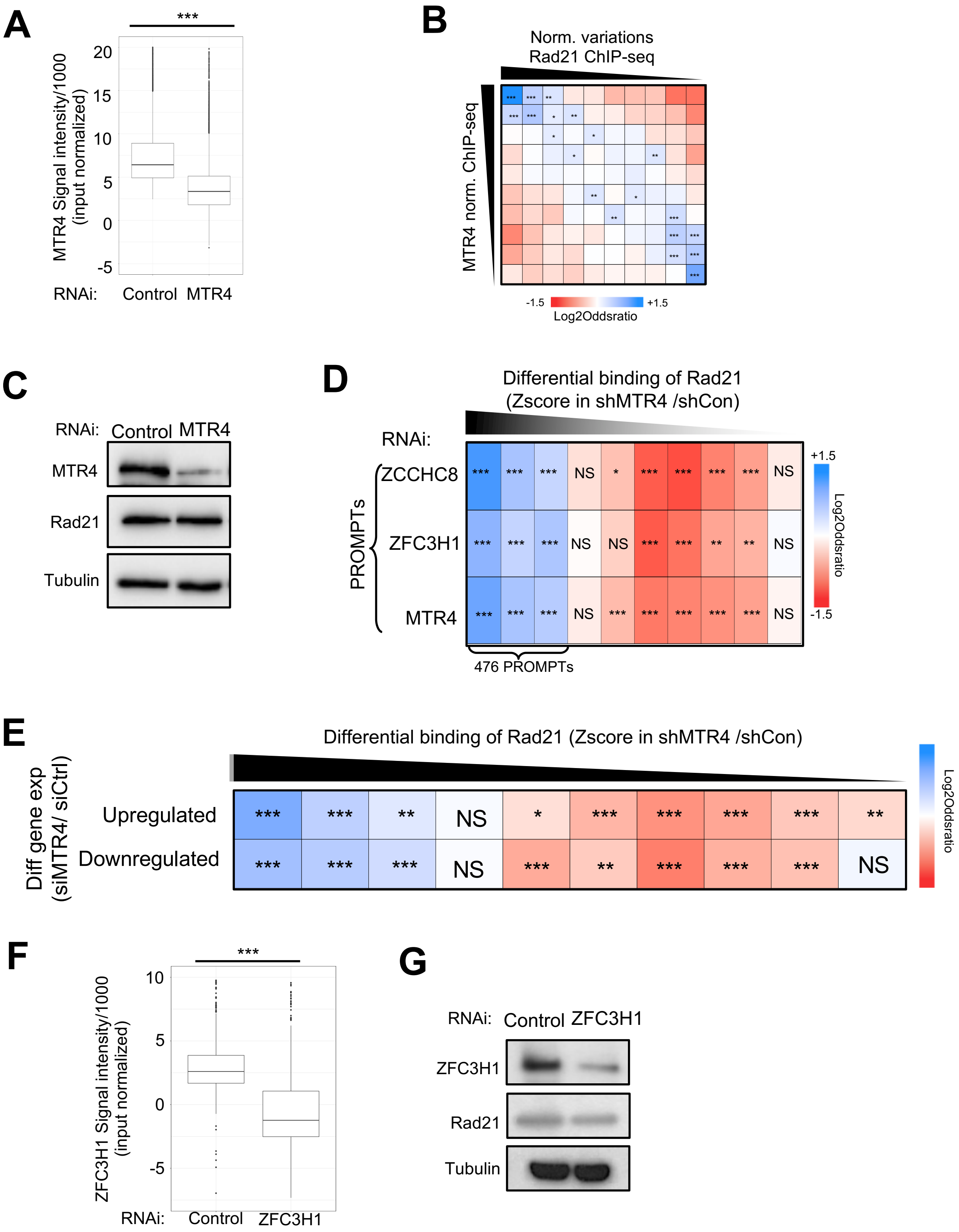


**Supplementary Fig. 5**

**Supplementary Fig. 5 Nuclear exosome regulates the association of cohesin with sites of long-range chromatin interactions.**

**A** Box plots of MTR4 ChIP-seq reads at regions bound by MTR4 in cells treated with shMTR4 or shControl, as indicated (*** $p < 0.001$, Wilcoxon test). **B** Intersection matrix showing Rad21 binding, stratified by the net variation in MTR4-depleted compared to control cells (horizontal axis), and the same sites stratified by MTR4 binding levels (vertical axis). The enrichment with distant H3K4me1 sites (not overlapping a TSS) was calculated using a Fisher exact test. **C** Whole cell extracts from HeLa cells treated with shRNA targeting MTR4 or a nontargeting control, as indicated, were analysed by Western blot using antibodies against Rad21 or tubulin. **D** Intersection matrix showing Rad21 binding, stratified by net variation upon MTR4 depletion compared to control, and increased or decreased gene expression, as indicated. The enrichment with PROMPTs was calculated using Fisher's exact test. **E** Box plots of ZFC3H1 ChIP-seq reads at regions bound by ZFC3H1 in cells treated with shZFC3H1 or shControl, as indicated (*** $p < 0.001$, Wilcoxon test). **F** Whole cell extracts from HeLa cells treated with shRNA targeting ZFC3H1 or a nontargeting control, as indicated, were analysed by Western blot using antibodies against Rad21, ZFC3H1 or tubulin.

**Table S1.** Antibodies used in this study.

| Antibody | Reference | Supplier |
| --- | --- | --- |
| MTR4 (ChIP-seq) | Ab70552 | Abcam |
| MTR4 (WB) | A300-614 | Bethyl laboratories |
| ZFC3H1 (ChIP-seq) | A301-457A | Bethyl laboratories |
| ZFC3H1 (WB) | A301-456A | Bethyl laboratories |
| ZCCHC8 (ChIP-seq) | A301-805A | Bethyl laboratories |
| ZCCHC8 (WB) | A301-806A | Bethyl laboratories |
| Rad21 (ChIP-seq and WB) | Ab992 | Abcam |
| TUBULIN | T9026 | Sigma-Aldrich |

**Table S2. Sequences of siRNAs used in this study.**

| siRNA | Sequence (5’ to 3’) |
| --- | --- |
| Control | ucugcaagguuaggcgucu (dTdT) |
| MTR4 | acacugagcuggaaaauaa(dTdT) |
